# Auditory brainstem responses to continuous natural speech in human listeners

**DOI:** 10.1101/192070

**Authors:** Ross K Maddox, Adrian KC Lee

## Abstract

Speech is an ecologically essential signal whose processing begins in the subcortical nuclei of the auditory brainstem, but there are few experimental options for studying these early responses under natural conditions. While encoding of continuous natural speech has been successfully probed in the cortex with neurophysiological tools such as electro- and magnetoencephalography, the rapidity of subcortical response components combined with unfavorable signal to noise ratios has prevented application of those methods to the brainstem. Instead, experiments have used thousands of repetitions of simple stimuli such as clicks, tonebursts, or brief spoken syllables, with deviations from those paradigms leading to ambiguity in the neural origins of measured responses. In this study we developed and tested a new way to measure the auditory brainstem response to ongoing, naturally uttered speech. We found a high degree of morphological similarity between the speech-evoked auditory brainstem responses (ABR) and the standard click-evoked ABR, notably a preserved wave V, the most prominent voltage peak in the standard click-evoked ABR. Because this method yields distinct peaks at latencies too short to originate from the cortex, the responses measured can be unambiguously determined to be subcortical in origin. The use of naturally uttered speech to evoke the ABR allows the design of engaging behavioral tasks, facilitating new investigations of the effects of cognitive processes like language processing and attention on brainstem processing.

**SIGNIFICANCE STATEMENT:** Speech processing is usually studied in the cortex, but it starts in the auditory brainstem. However, a paradigm for studying brainstem processing of continuous natural speech in human listeners has been elusive due to practical limitations. Here we adapt methods that have been employed for studying cortical activity to the auditory brainstem. We measure the response to continuous natural speech and show that it is highly similar to the click-evoked response. The method also allows simultaneous investigation of cortical activity with no added recording time. This discovery paves the way for studies of speech processing in the human brainstem, including its interactions with higher order cognitive processes originating in the cortex.

## INTRODUCTION

When speech enters the ear and is encoded by the cochlea, it goes on to be processed by an ascending pathway that spans the auditory nerve, brainstem, and thalamus before reaching the cortex. Far from being relays, these subcortical nuclei perform a dazzling array of important functions, from sound localization (Grothe and Pecka, 2014) to vowel coding (Carney et al., 2015), making their function essential to understand. In humans, the primary method for measuring activity in subcortical nuclei is the auditory brainstem response (ABR): a highly stereotyped scalp potential in the first ∼10 ms following a very brief stimulus such as a click, recorded through electroencephalography (EEG) (Burkard et al., 2006). The potential comprises components referred to as waves, given Roman numerals I–VII according to their latency. Individual waves have been tied to activity in specific parts of the ascending pathway: wave I (∼2 ms latency) is driven by auditory nerve activity, wave III (∼4 ms) by the cochlear nucleus, and wave V (∼6 ms) principally by the lateral lemniscus (Møller et al., 1995). However, because the waves are so rapid, and the signal-to-noise ratio so low, the ABR must be measured by presenting thousands of repeated punctate stimuli. Thus, while there are important neuroscience questions regarding how subcortical nuclei process natural stimuli like speech, or how they might be affected by cognitive processes through efferent feedback (Terreros and Delano, 2015), the practical limitations of the ABR paradigm make it primarily a clinical tool.

One common method for measuring the brainstem response to speech is the complex ABR (cABR) (Skoe and Kraus, 2010). The cABR represents the averaged response to repetitions of a short spoken syllable (e.g., a ∼40 ms “da”). It can be analyzed in the time domain, but because the stimulus is longer than the response, ambiguity about the origin of response components arises. The voiced part of the speech elicits a frequency following response (FFR) that can be analyzed in the frequency domain. The FFR at the stimulus’s harmonics is reasoned to have subcortical origins because of the lower frequency phase-locking limit in the auditory cortex (Joris et al., 2004), but a recent magnetoencephalography study showed cortical contributions to the FFR (Coffey et al., 2016), rendering strong conclusions about exclusively subcortical phenomena difficult to make.

A different method, used for studying cortical activity, treats the auditory evoked potential as the impulse response of a linear system, which can be mathematically derived from known input and output signals (Aiken and Picton, 2008; Lalor et al., 2009; Lalor and Foxe, 2010; Ding and Simon, 2012a, 2012b). Continuous natural speech is presented (input) while EEG is recorded (output), and the brain’s response is calculated through linear regression. Rather than raw audio, the regressor (i.e., input) used is the amplitude envelope, which by construction contains no fast fluctuations, making it too slow for studying subcortical nuclei. A recent study aimed at the brainstem used the amplitude envelope of a speech stimulus’s fundamental frequency as input, and the envelope of the EEG at that frequency as output (Reichenbach et al., 2016). The response is a single wave with a peak latency of 10 ms, suggesting brainstem involvement, but a width of 100 ms, making it impossible to exclude cortical contributions.

Here we measured auditory brainstem activity in response to natural speech using a new paradigm. The methods were based on cortical studies, with an important difference: the regressor was the rectified speech audio, meaning that fine structure was largely preserved. The speech-evoked responses were very similar to click-evoked ABRs, most notably in the presence of a distinct wave V. Because the latency of the wave V peak is shorter than a cortical source could produce, it can be unambiguously attributed to subcortical generators. We show that it is possible to study speech processing in the human brainstem, paving the way for subcortical studies of attention, language, and other cognitive processes.

## MATERIALS AND METHODS

### Experimental design and statistical analysis

Our goal was to measure the speech-evoked ABR in human listeners and validate it against the click-evoked response. We first recorded click-evoked responses to psuedorandomly timed click trains and then validated them against the responses evoked by standard, periodic click trains. We then compared the speech-evoked response to the pseudorandom click-evoked response. We validated by comparing the overall morphology, as well as the presence and latency of wave V in the speech-evoked response.

All subjects’ click- and speech-evoked responses were plotted individually. To compare the similarity of two responses from a single subject (e.g., the click-evoked response to the speech-evoked response), Pearson’s product-moment correlation was used. The median and interquartile range of each distribution of correlation coefficients across subjects was reported, in addition to plotting its histogram. Two distributions of correlation coefficients were compared using Wilcoxon’s signed-rank test for non-normal distributions. When comparing wave V latencies across stimulus conditions, a paired Student’s t-test was used to determine if the means differed.

### Subjects

All experiments were done under a protocol approved by the University of Washington Institutional Review Board. All subjects gave informed consent prior to participation, and were compensated for their time. We collected data from 24 subjects (17 females). The mean age was 27.8 years, with a standard deviation of 6.9 and a range of 19–45. Subjects had normal hearing, defined as audiometric thresholds of 20 dB HL or better in both ears at octave frequencies ranging from 250 to 8000 Hz. All subjects identified English as their first language except for two, who identified a different language but had been speaking English daily for over twenty years.

### EEG recording

Scalp potentials were recorded with passive Ag/AgCl electrodes, with the positive and negative electrodes connected to a differential preamplifier (Brainvision LLC, Greenboro, SC). The positive electrode was at location FCz in standard 10-20 coordinate system. The negative (reference) electrode was clipped onto the subject’s left earlobe. The ground electrode was placed at Fpz. Data were high-passed at 0.1 Hz during recording (additional filtering occurred offline).

Subjects were seated in a comfortable chair in a sound-treated room (IAC, North Aurora, IL). They were not asked to attend the stimuli. Instead, they faced a computer monitor showing silent episodes of “Shaun the Sheep” (Aardman Animations, 2007), an animated show that has no talking, making subtitles unnecessary. They were first presented with 40 epochs of speech stimuli for calculating the speech ABR, and then were presented with 10 minutes of click stimuli (twenty repetitions of a frozen 30 s epoch). All stimuli were presented over insert earphones (ER-2, Etymotic Research, Elk Grove, IL) which were plugged into a stimulus presentation system consisting of a real-time processor and a headphone amplifier (RP2.1 and HB7, respectively, Tucker Davis Technologies, Alachua, FL). Stimulus presentation was controlled with a python script using publicly available software (available at https://github.com/LABSN/expyfun).

### Speech stimuli

Speech stimuli were taken from two audiobooks. The first was *A Wrinkle in Time* (L’Engle, 2012), read by a female narrator. The second was *The Alchemyst* (Scott, 2007), read by a male narrator. The audiobooks were purchased on compact disc and ripped to uncompressed wav files to avoid data compression artifacts. They were resampled to 24,414 Hz, the native rate of the RP2 presentation system. They were then processed so that any silent pauses in the speech longer than 0.5 s were truncated to 0.5 s. Because the ABR is principally driven by higher stimulus frequencies (Abdala and Folsom, 1995), the speech was gently high-passed with a first-order Butterworth filter with a cutoff of 1,000 Hz and a slope of 6 dB / octave. The speech was still natural sounding and intelligible. This filter also helped to compensate for low-frequency spectral differences between the male and female narrator around their fundamental frequencies. After that, the speech was normalized to an average root-mean-square amplitude that matched that of a 1 kHz tone at 75 dB SPL. Figures 1A,D,G show the pressure waveform of the word “Thursday” spoken by the male narrator, the spectrogram of that word’s first syllable, and the power spectral density (PSD) of a 30 s segment of the female and male speech stimuli.

**Figure 1.**
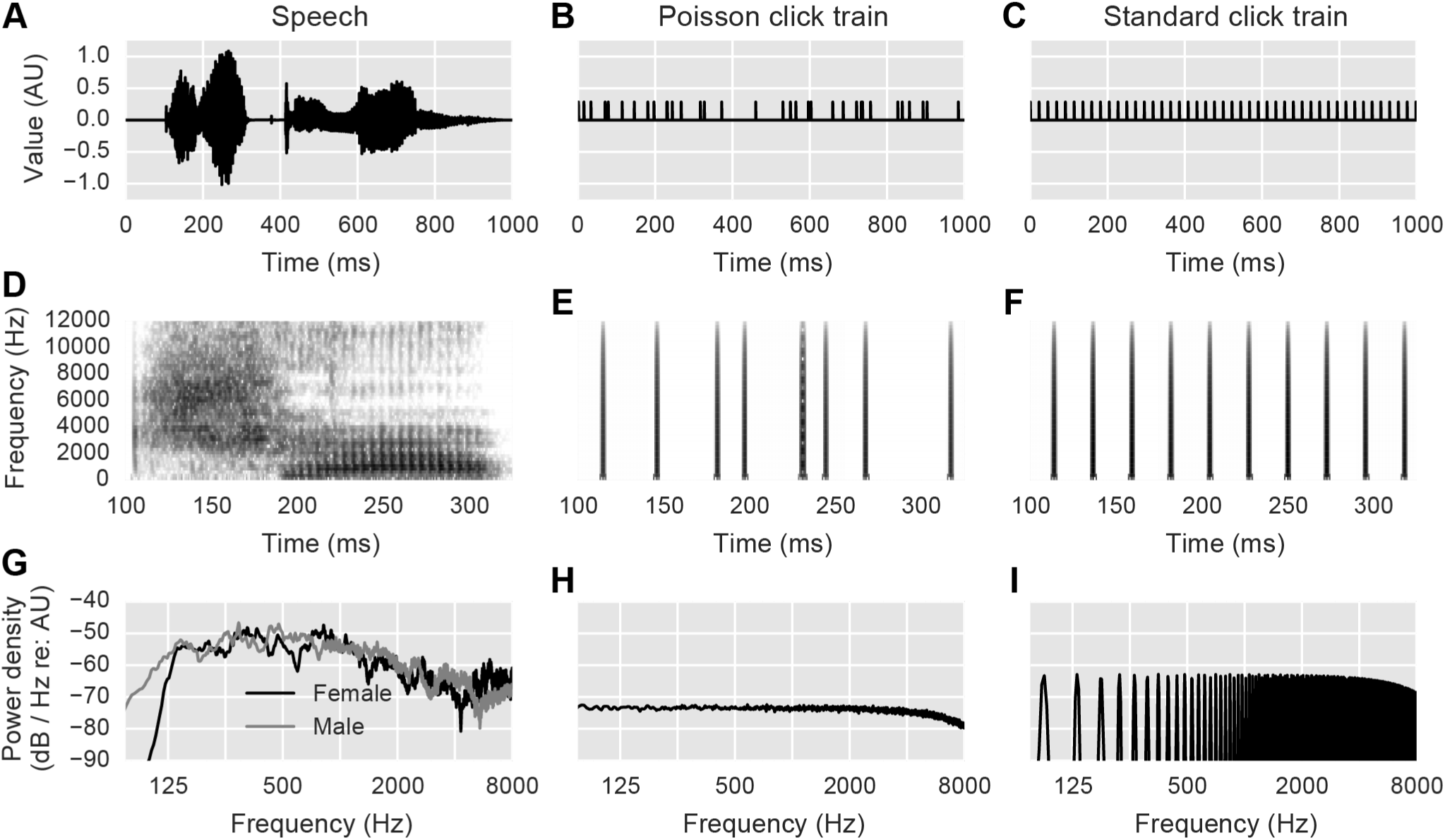
Acoustic stimuli. (A,B,C) Pressure waveforms for one second of speech, Poisson click train, and standard periodic click train, respectively. Vertical scale is arbitrary but consistent across plots. (D,E,F) Spectrograms of a smaller excerpt of the above stimuli, with darker colors corresponding to higher power. (G,H,I) Power spectral density plots of the above stimuli, calculated from 30 s of data using Welch’s method with a segment length of 5.67 ms, segment overlap of 50%, and Hann window.

The audiobooks were then sectioned into epochs of 64 s, including a 1 s raised cosine fade-in and fade-out. The last four seconds of each epoch were repeated as the first four seconds of the next one, so that subjects could pick up where they left off in the story (if they were listening), meaning that 60 s of novel speech was presented in each epoch. The stimuli were not new to the subjects—before this passive listening task, they had completed a session using the same stimuli where they had to answer questions about the content they had just heard. Data from that task were for a different scientific question and do not appear here. These minute-long excerpts were presented in sequence, two from one story and then in alternating sets of four, finishing with two epochs from the second story. Speech stimuli were presented diotically.

### Click stimuli

Click stimuli were aperiodic trains of rarefaction clicks lasting 82 μs (representing two samples at the 24,414 Hz sampling rate, which was closest possible to the standard 100 μs click duration with our hardware). Clicks were timed according to a Poisson point process with a rate of 44.1 clicks / s. The timing of one click had no correlation with the timing of any other click in the train, rendering the sequence spectrally white in the statistical sense. A pair of 30 s sequences was created and presented dichotically 20 times to each subject, meaning that 26,460 clicks contributed to each ear’s response. The responses presented herein are the sum of the monaural responses. Clicks were presented at 75 dB peak-to-peak equivalent SPL (i.e., the amplitude of clicks matched the peak-to-peak amplitude of a 1 kHz sinusoid presented at 75 dB SPL).

While no previous study has used exactly this type of click timing, several have used various types of pseudorandom sequences (Burkard et al., 1990; Thornton and Slaven, 1993; Delgado and Ozdamar, 2004; Holt and Özdamar, 2014). Uniformly, these studies find that the ABRs from randomized versus periodic click trains are highly similar at the same stimulation rates. Random timing has two main benefits over the much more common periodic timing: 1) the analysis window for the response can be extended arbitrarily to any beginning and end point without fear of temporal wrapping, and 2) no high-pass filtering is necessary to remove the strong frequency component at the (periodic) presentation rate, because it does not exist. A third benefit, specific to this study, is that the same analysis could be done to compute the speech-evoked and the click-evoked ABR, yielding a more direct comparison between the two. Figures 1B,E,H show part of a Poisson click train in the same manner that Figs. 1A,D,G do for speech.

To be sure that the click paradigm we used yielded results matching standard ABRs evoked with periodic click trains, we also collected ABRs using periodic click trains of the same rate of 44.1 clicks / s, presented diotically. Periodic trains were also presented in twenty epochs of 30 s, yielding the same total sweep count of 26,460. The periodic click train stimulus is shown in Figs. 1C,F,I.

### Data analysis

Responses to both speech and click train stimuli were found through deconvolution, in a manner broadly similar to previous papers focused on cortical activity (Lalor et al., 2009; Lalor and Foxe, 2010). The essence of deconvolution is determining the impulse response of a linear time-invariant system given a known input (here, the processed continuous speech signal) and a known output (here, the recorded scalp potential). The methods in this study vary from previous ones in the preprocessing steps, but otherwise utilize essentially the same mathematical principles.

#### Speech stimuli preprocessing

Before we could derive the speech response, we needed to calculate the regressor from the audio data. The auditory brain is mostly agnostic to the sign of an acoustic input, as evidenced by the high degree of similarity between evoked responses to compression versus rarefaction clicks (Møller et al., 1995). For this reason, some sort of rectifying nonlinearity applied to the input speech is needed as a preprocessing step. We used half-wave rectification. Specifically, we performed all analyses twice— once keeping the positive peaks, and then a second time keeping the inverted negative peaks—and then averaged the resulting responses, in a process akin to the compound peristimulus time histogram used by Pfeiffer and Kim (1972). This significantly reduced, but did not eliminate, stimulus artifacts, similar to the common technique of alternating polarity in the click-evoked ABR (Hall III, 2006). Following rectification, the data were downsampled from 24,414 Hz to the EEG recording rate of 10,000 Hz.

#### Click train preprocessing

Owing to its extreme sparsity, downsampling a click train using standard methods would result in significant signal processing artifact, viz., Gibbs ringing. We instead used the list of click times from the original click train (24,414 Hz sampling rate) and created a click train at 10,000 Hz sampling rate by placing unit-height single-sample impulses at the closest integer indices corresponding the original click times.

When the input to a system has a white power spectrum, the system’s impulse response can be determined as the cross-correlation of the input and output. For a click train, which is essentially a series of unit-height single-sample impulses, the deconvolved impulse response becomes equivalent to the click-triggered average, which is how ABRs are usually calculated. This results in a convenient parity between the typical averaging methods used for ABR and the deconvolution used here. In other words: rather than using a completely new mode of analysis for ABR (deconvolution), we have instead generalized the methods already in use to be appropriate for arbitrary stimuli, beyond click trains.

#### EEG preprocessing

EEG data were first high-pass filtered at 1 Hz (first-order Butterworth), and then notch filtered at 60, 180, and 300 Hz with 5 Hz wide second-order infinite impulse response notch filters, designed with the *iirnotch* function of the SciPy python package (RRID:SCR_008058). Because of the continuous nature of the stimuli, no epoch rejection was done. Instead, any time the EEG potential crossed ±100 μV, a 1 s segment of the response was zeroed, centered around the offending sample, removing it from the calculation. This operation effectively reduced the energy of an epoch. So that the amplitude of the calculated response was not affected, the EEG data for each epoch was multiplied by a corrective gain factor *g*_*r*_:

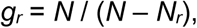

where *N* is the total number of samples in the epoch and *N_r_* is the number of rejected samples. After filtering and resampling, the data were segmented into epochs that started with the stimulus onset and ended 100 ms after the stimulus (epochs were thus 64.1 s long for speech stimuli and 30.1 s long for clicks).

#### Response calculation

We used linear least-squares regression to calculate the responses, as in previous work (Lalor et al., 2009). The response was considered to be the weights over a range of time lags that best approximated the EEG output as the weighted sum of the input stimulus regressor over those lags. For the sake of computational and memory efficiency, the stimulus autocorrelation matrix and stimulus-response cross-correlation were both calculated via their Fourier counterparts using frequency-domain multiplication. These specific methods have been incorporated into the mne-python package (Gramfort et al., 2013) (RRID:SCR_005972) (https://github.com/mne-tools/mne-python/blob/8fc2a545f494de0f828b931f2285dbff426e72ad/mne/decoding/time_delaying_ridge.py). No regularization was employed. The response weights were calculated over the range of lags spanning -150 to 350 ms. After the response was calculated, it was low-pass filtered at 2,000 Hz (first-order Butterworth), and then baseline corrected by subtracting the mean potential between -10 and 0 ms from the whole response. For the speech stimuli, the response to each narrator was calculated separately, and then averaged to calculate each subject’s speech-evoked response.

#### Speech-evoked response amplitude normalization

Auditory onsets elicit much larger responses than ongoing stimulus energy due to adaptation (Thornton and Slaven, 1993). However, this non-linear adaptation is not accounted for by the linear regression. For that reason, the raw speech-evoked responses, for which the majority of the stimulus energy can be considered “ongoing,” were much smaller than the click-evoked responses, whose stimuli are essentially a series of onsets. To correct for this, we computed a single empirical subject-specific normalization factor, *g*_*n*_, that put the speech-evoked responses in a similar amplitude range as the click-evoked ones:

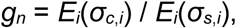

where *σ*_*c,i*_ is the standard deviation of subject *i*’s click-evoked response in the range of 0–30 ms, *σ*_*s*,*i*_ is the same for the speech-evoked response, and *E*_*i*_ represents the mean over subjects. All speech-evoked responses shown in microvolts have been multiplied by *g*_*n*_. In our study *g*_*n*_ had a value of 27.5, but it must be stressed that this value depends on the unitless scale chosen for storing the digital audio, and is thus not suitable for use in other studies. For this reason no direct amplitude comparisons were made between click- and speech-evoked responses. Instead, their morphologies and wave V latencies were compared.

### Standard ABR measurement

The ABR to the periodic click trains was calculated through traditional averaging rather than regression. The raw data were notch filtered to remove line noise and low-pass filtered at 2,000 Hz as described above. However, the high-pass filter was different: a causal second order Butterworth filter with a cutoff of 150 Hz was used to be consistent with standard practice and to generate a canonical waveform (Burkard et al., 2006; Hall III, 2006). The response to each click presentation was then epoched from -3 ms to 19.7 ms, which was the longest window allowed by the periodic click rate of 44.1 clicks / s before temporal wrapping occurred. Filtered epochs were rejected if the peak-to-peak amplitude exceeded 100 μV.

## RESULTS

### Poisson click trains yield canonical ABRs

Responses to Poisson click trains were used as the benchmark to which the speech-evoked responses were compared. Even though similar types of pseudorandom stimuli have been used in the past, it was important to confirm that these specific stimuli used here provided canonical ABR waveforms. The grand average periodic and Poisson click trains are shown overlaid in Fig. 2A (both shown high-pass filtered at 150 Hz). To quantify their similarity, we computed Pearson’s correlation coefficient between the two waveforms for each subject between lags of 0 and 19.7 ms. The median correlation was 0.89 (interquartile range 0.82–0.92), indicating a very high degree of similarity. The histogram of correlations is shown in Fig. 2B.

**Figure 2.**
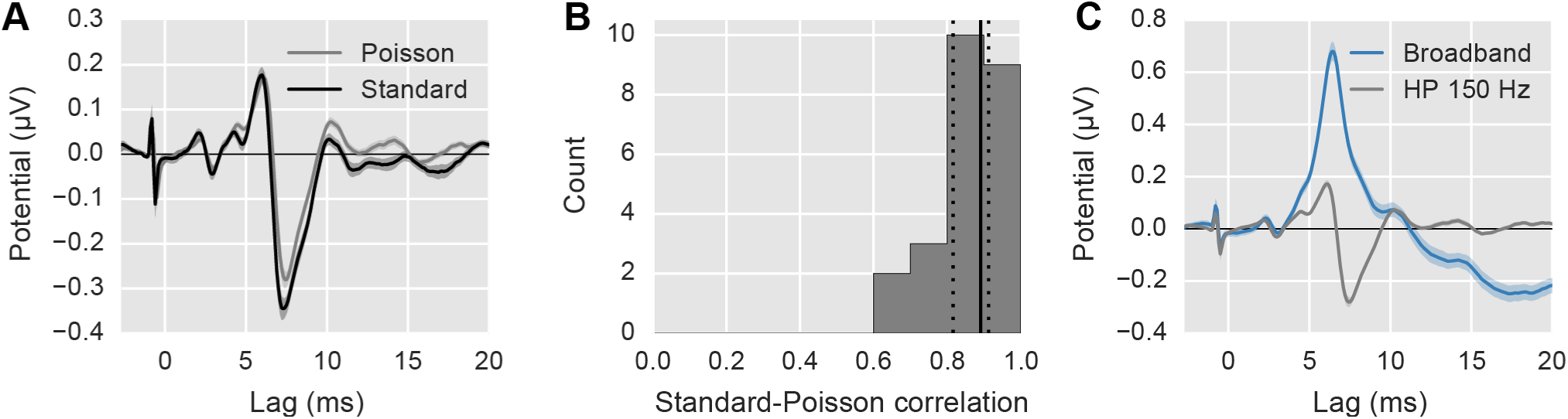
Comparison of ABR to standard periodic click trains and Poisson click trains. (A) The average ABR waveform evoked by the standard, periodic click train at 44.1 clicks / s (black) and the psudorandom Poisson click train (gray; 44.1 clicks / s overall rate). Areas show ±1 SEM. Both responses are high-pass filtered at 150 Hz. The spike at -1 ms is a stimulus artifact, and occurs before 0 ms to compensate for the 1 ms tube delay of the earphones. (B) The histogram of correlation coefficients between the standard and Poisson click-evoked ABRs. Solid/dotted black lines show median/quartiles. (C) Comparison of the Poisson click-evoked ABR with 150 Hz high-pass filtering (gray) and without (i.e., broadband; blue). The latter is used as the benchmark response for the remainder of the study.

Figure 2C shows the average Poisson click-evoked response under two filtering conditions: 1) high-pass filtered at 150 Hz as in Fig. 2A, and 2) broadband (high-passed at 1 Hz as described in the EEG pre-processing methods section above). The latter will be used henceforth as the click-evoked ABR to which the speech-evoked ABR is compared. It is thus important to note that even though these responses seem to have morphological differences from the “standard” ABR, that is simply because using pseudorandom click timing obviates the need for high-pass filtering, and that filtering was bypassed in the interest of comparing the whole responses. The wideband responses we obtained here using Poisson click trains were highly similar in shape, amplitude, and latency to previous wideband (5 Hz high-pass) ABRs obtained using low rate (11 Hz) periodic clicks (Gu et al., 2012), and were much more efficient to obtain.

### Early speech-evoked responses exhibit brainstem response characteristics

Broadly speaking, there were strong similarities between the early (< 30 ms) click-evoked and speech-evoked responses (Fig. 3A). In this latency range, responses are likely to progress from brainstem to thalamus and primary auditory cortex as latency increases. We will first make whole-waveform comparisons, and then consider specific canonical ABR components.

**Figure 3.**
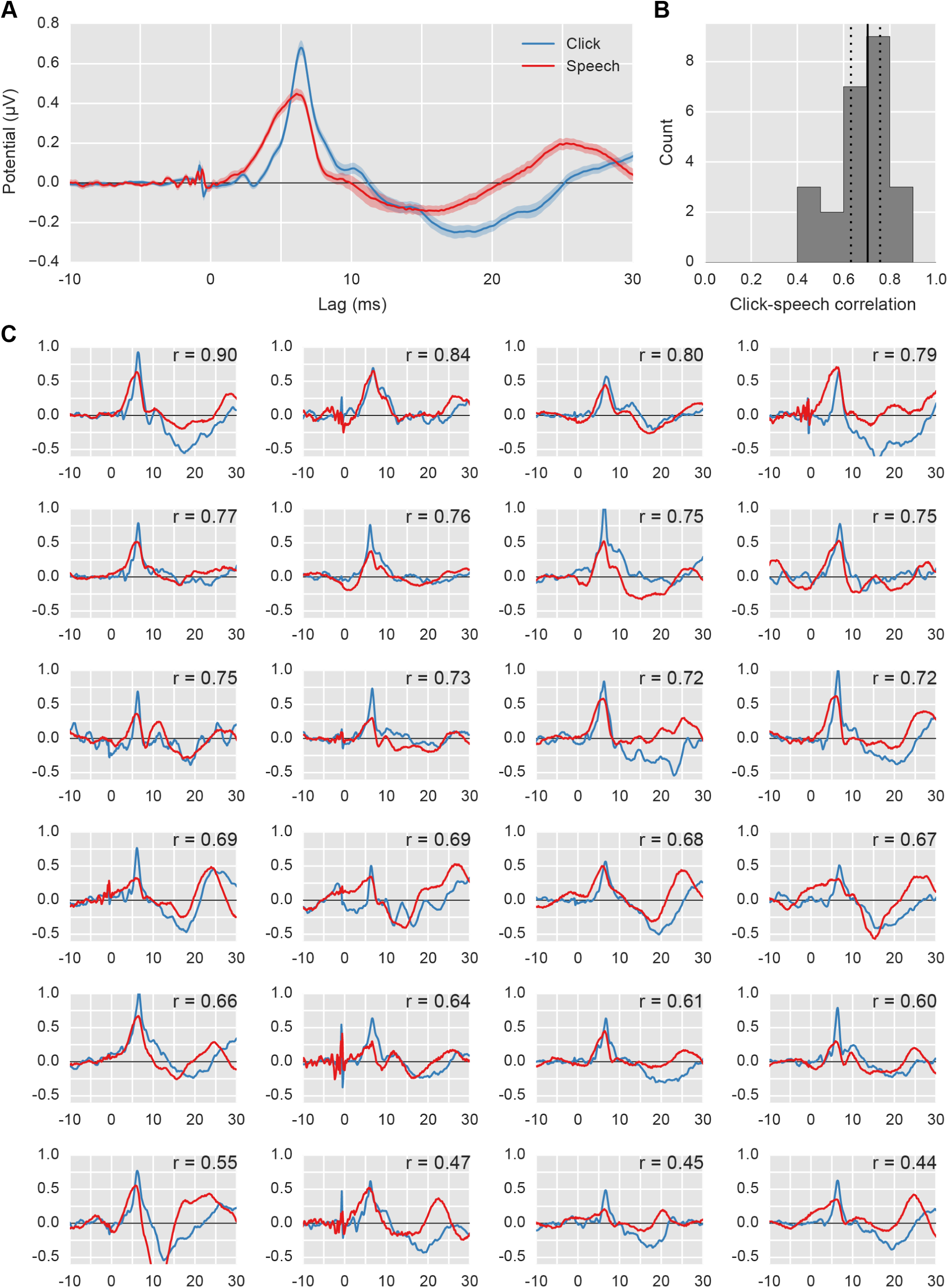
Comparison of click responses (blue) with speech responses (red). (A) The average waveform across subjects (areas show ±1 SEM). (B) The histogram of correlation coefficients between the click-evoked and speech-evoked stimuli for each subject. Solid/dotted black lines show median/quartiles. (C) Individual subject responses. The correlation coefficient is shown in the upper right corner.

To compare the overall waveforms, we computed Pearson’s correlation coefficient of the speech- and click-evoked waveforms for each subject in the range of 0–30 ms (Fig. 3B). The median correlation coefficient was 0.70 (interquartile range 0.63–0.75). Figure 3C shows each subject’s click- and speech-evoked response, in descending correlation order. In our speech-evoked responses, waves I–IV were “smeared” together. However, we found a clear wave V in individual subjects’ responses as well as the grand average. Wave VI was also visible in the grand average, but was less consistent at the individual-subject level.

We identified wave V by low-pass filtering at 1,000 Hz with a zero-phase filter and finding the peak of the waveform in the 5–7 ms range. For the click-evoked responses, wave V was present for all subjects, with a latency of 6.50 ± 0.25 ms (mean ± standard deviation). For speech-evoked responses, wave V was present for all subjects, with a latency of 6.17 ± 0.30 ms. The speech-evoked wave V preceded the click-evoked by 0.26 ms (t(23) = 6.6, p = 1×10^−6^, paired t-test). As shown in Fig. 4, the click-evoked and speech-evoked wave V latencies were correlated across subjects (r = 0.75, p = 3×10^−5^, Pearson’s product-moment). This shows a strong correspondence between the click-evoked and speech-evoked ABR.

**Figure 4.**
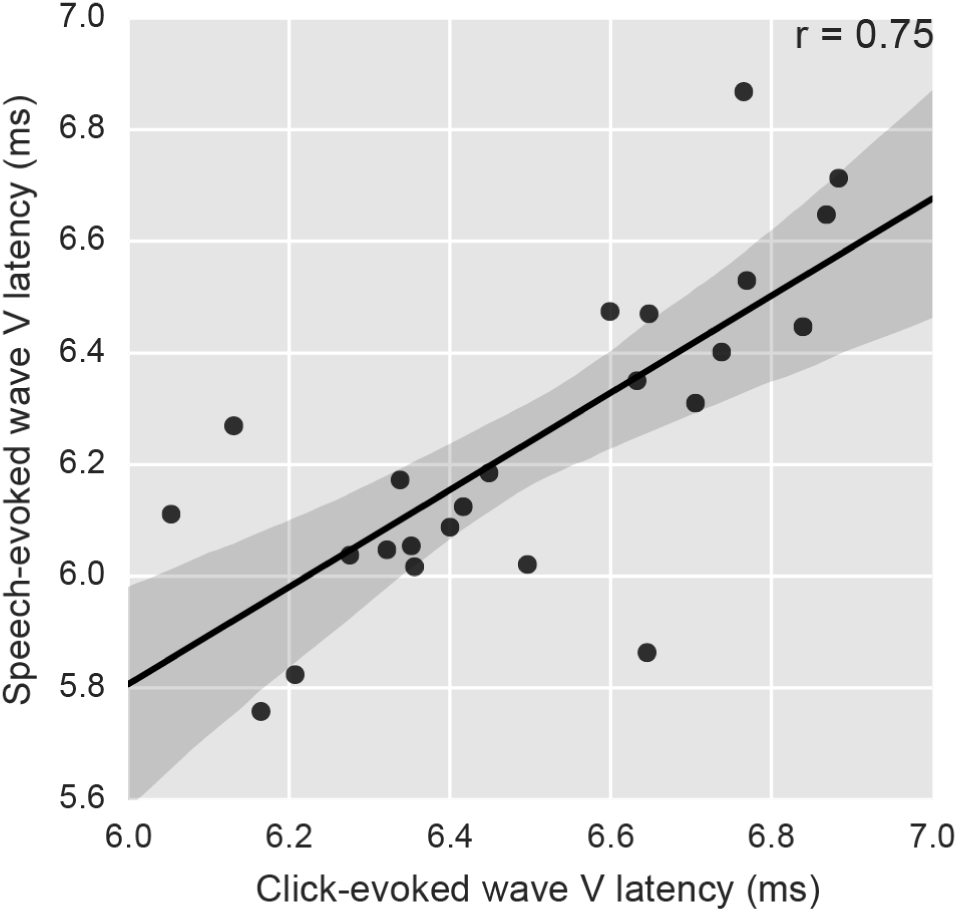
Speech-evoked versus click-evoked wave V latencies across subjects. The strong correlation across subjects points to common neural generators. Points have been jittered slightly to prevent overlap. Regression line is shown with the 95% confidence interval shaded.

In some subjects’ speech-evoked waveforms there are early peaks that seem to resemble waves I and III. However, these are likely driven by recording artifacts (electromagnetic leakage of the earphone driving signal into the EEG electrode recording). While it may have been possible to reduce these artifacts further through additional signal processing, we did not do that for sake of simplicity and transparency. However, it is important to note that a simple modification to the paradigm—alternating the polarity of the speech stimulus—should all but remove stimulus artifacts in the future. This could be done at the level of the 64 s epochs, or it could be done at the word or phrase level, as long as the phase inversions were hidden by silent gaps in the speech.

### Speech responses depend minimally on sex of talker stimuli

One important question is whether the speech-evoked response maintains its morphology independent of the specific input stimulus, or if it depends on the specific narrator. To investigate this, we split the responses to male- and female-narrated trials and compared them to determine the role that the difference in the narrators’ input spectra might play. The grand average waveforms for the two narrators are of the same magnitude and overall shape, despite the differing spectra of their input stimuli (Fig. 5A). The median female-male correlation coefficient was 0.73 (interquartile range 0.60–0.83; Fig. 5B). Figure 5C shows each subject’s response to the female- and male-narrated speech, in the same order as Fig. 3C to allow comparison.

**Figure 5.**
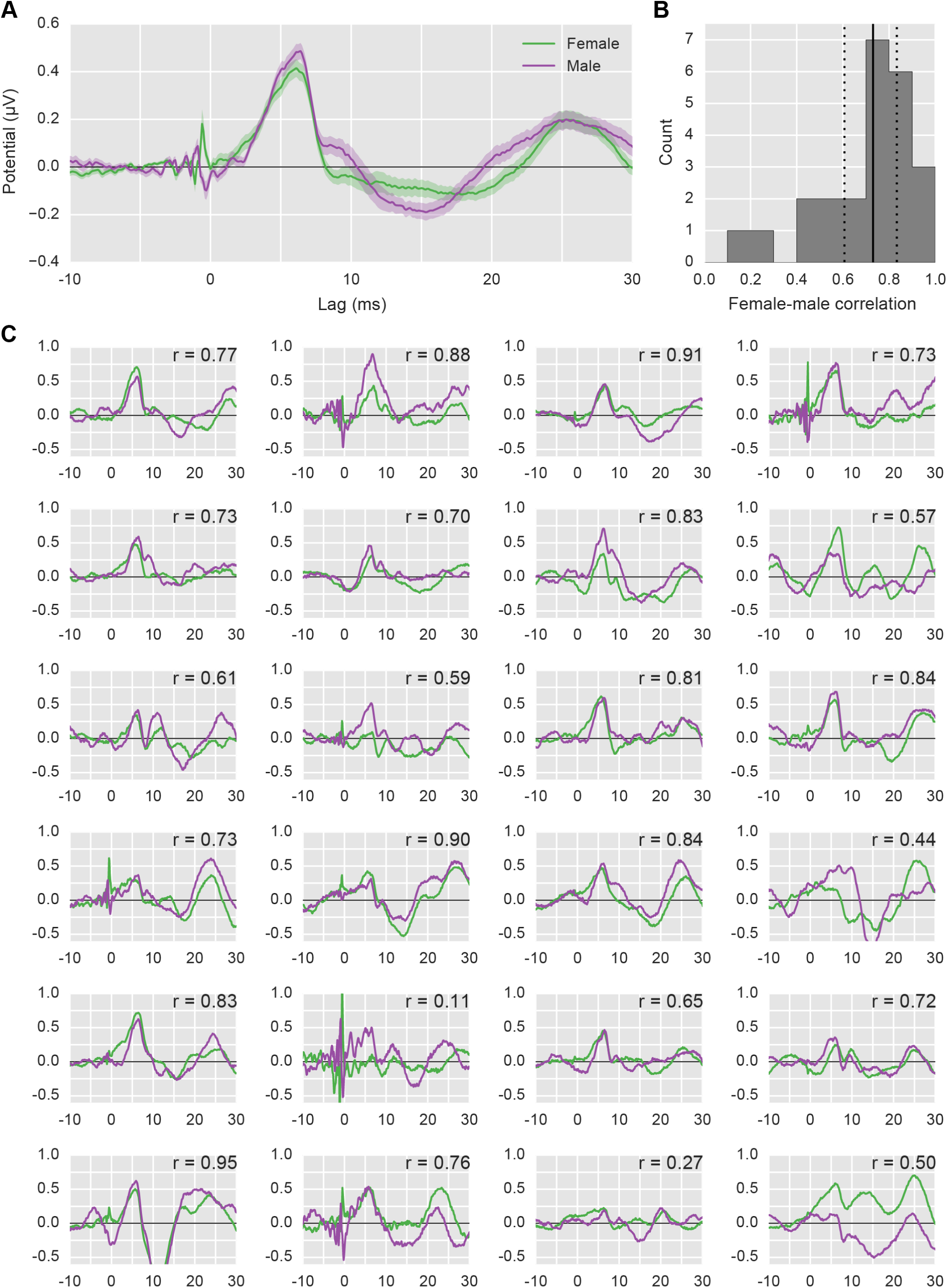
Comparison of female-narrated responses (green) with male-narrated responses (purple). (A) The average waveform across subjects (areas show ±1 SEM). (B) The histogram of correlation coefficients between the female-evoked and male-evoked stimuli for each subject. Solid/dotted black lines show median/quartiles. (C) Individual subject responses arranged in the same order as Fig. 3C for easy comparison. The correlation coefficient is shown in the upper right corner. The poor correlation of the worst subject (r = 0.11) is the result of a strong stimulus artifact.

While perfect overlap would be indicated by correlation coefficients of 1.0, splitting the data in half (viz., into male- and female-narrated epochs) adds noise to each of the responses. To put the male-female correlation coefficients in context, we can split the data a different way and compare. We split the data into halves that contained the same number of male and female epochs (i.e., each split contained 10 male and 10 female trials). We then compared those waveforms in the same way as above. The median correlation coefficient between splits was 0.83 (interquartile range 0.70–0.91). We compared the male-female split coefficients to these arbitrarily split coefficients, and found a significant difference (T(23) = 58, p = 0.009, Wilcoxon signed-rank test). This indicates that while the responses to female and male-uttered speech are very similar, there is still some dependence on the stimulus.

## DISCUSSION

### Early speech responses are interpretable as ABRs

The major goal of this work was to study the response of the human auditory brainstem to naturally spoken, continuous speech. We derived the speech-evoked responses using regression and validated them against click-evoked responses. Comparison of the speech-evoked and click-evoked ABR revealed a high degree of morphological similarity between waveforms, similar overall wave V latencies, and a strong correlation between click- and speech-evoked wave V latency across subjects. Taken together, these results show that the speech-evoked ABR is just that—the response of the auditory brainstem.

Incoming acoustic information travels up the auditory pathway in an initial feedforward sweep, from brainstem to thalamus to cortex. Because the response calculated here is broadband, distinct components over the range of latencies were preserved. We can thus “localize through latency” and logically conclude that the peak in the response at 6 ms has subcortical origins, because it is too soon after the stimulus to be cortical, where the earliest estimated latencies are 11–14 ms (Wassenhove and Schroeder, 2012). This eschews the problem of source mixing when attempting to determine brainstem activity through spatial means, such as beamforming and dipole fits. However, as discussed below, our method does not preclude those analyses—rather it complements them and facilitates their use, particularly at longer latencies where sources have cortical origins more appropriate for spatial filtering.

### Subcortical and cortical responses are available simultaneously

While the focus of this work is on the brainstem and midbrain responses, these methods can be used to measure both subcortical and cortical activity. Simultaneous subcortical and cortical measurements are possible with the cABR (Skoe and Kraus, 2010), but the differing parameters for number of trials and inter-stimulus interval needed mean that recording paradigms can be very long. Work aimed at optimal parameters for simultaneous subcortical-cortical recordings has been successful (Bidelman, 2015), but still necessarily results in compromises. The present methods allow simultaneous measurement with no additional recording time and no limitations on the response window due to inter-stimulus interval.

This flexibility is illustrated in Fig. 6. Figure 6A shows the speech-evoked ABR, Fig. 6B extends the window and employs a low-pass filter appropriate for viewing the middle latency response (Hall III, 2006), and Fig. 6C extends the time window further and lowers the low-pass frequency to accentuate late auditory evoked potentials of cortical origin. If a full electrode montage (and sufficient hard drive space) is available, the interaction of brainstem processing with any number of cortical processes is now possible to investigate under natural conditions.

**Figure 6.**
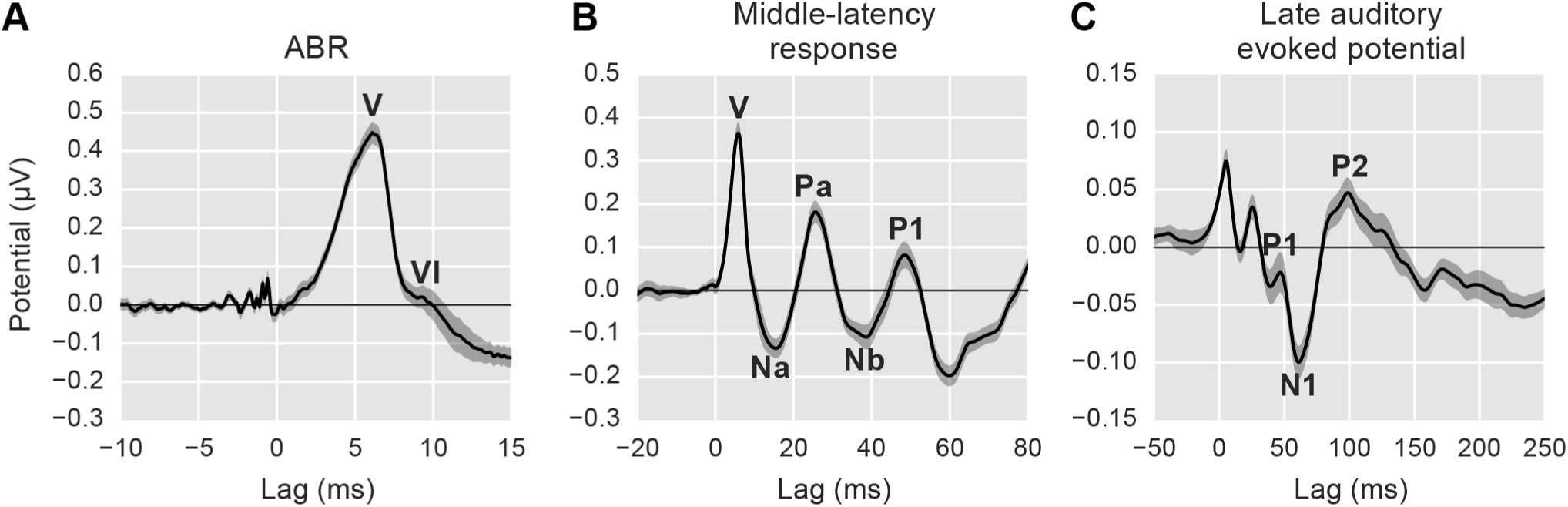
Changes to the range of lags and filtering parameters allows early, middle, and late responses to be analyzed from the same recording. (A) The speech-evoked auditory brainstem response with canonical waves V and VI labeled. (B) The middle latency response with its canonical waves labeled (low-pass frequency: 200 Hz). (C) The late auditory evoked potential with its canonical waves labeled (low-pass frequency: 20 Hz). Shaded areas show ±1 SEM.

### Filtering must be done carefully

It is common practice in EEG experiments to use zero-phase filters whose impulse responses are non-causal and symmetric about 0 lag. This is done to preserve the latencies of the peaks and is appropriate in most cases. However, the strength of the present approach lies in using the latency of the response peaks to confirm their subcortical origin. If a non-causal filter is used to filter the EEG data, then it is possible that a peak at a latency corresponding to cortical activity (e.g., 25 ms) could affect the response waveform at brainstem latencies (e.g., 6 ms). This could have the result of erroneous findings that attribute cortical phenomena to subcortical nuclei. Thus, the following two guidelines should be followed for experiments specifically aimed at the auditory brainstem. First, EEG data should be filtered with causal filters. Second, when calculating regressors, any filtering that is done to the input stimulus should be anti-causal (i.e., with an impulse response has values only at negative lags). The latter can be practically accomplished by reversing the signal in time, filtering it with a standard causal filter, and then reversing that result. Using causal filters will inevitably affect the latencies of peaks, but this can be mitigated by filtering sparingly (i.e., as broadband as the specific analyses will allow) with low-order filters.

### Responses to arbitrary stimuli can be measured

For a spectrally rich but non-white stimulus like speech, an important step in deconvolution is whitening the input stimulus. For a linear system, two broadband stimuli with different spectra should yield the same impulse response. However, there is no such guarantee for a non-linear system like the auditory system.

The present study suggests that a range of stimuli can be used. First, we consider the main comparison: speech-evoked to click-evoked ABR. Natural speech is different by almost any metric from Poisson click trains, and yet the responses that we find through regression are very similar (Fig. 3A,B). Second, we consider the responses to female versus male speech. Males typically speak at a fundamental frequency about half that of females. Such a difference, when estimating the response of a highly non-linear system using linear methods, could have resulted in major differences in the response waveforms, but this was not the case (Fig. 5A,B). Taken together, it is reasonable to expect that the technique could be applied to other real-world non-speech stimuli such as music or environmental sounds, as well any spectrally rich synthetic stimulus of interest in the lab.

Despite the similarity between responses to different stimuli, the differences (e.g. between the female and male speech-evoked responses) represent a caveat. In future studies, experimenters must be careful in making comparisons between responses across conditions that did not use identical stimuli. We suggest that these methods will be most useful in cases where the acoustic stimuli can be counterbalanced across conditions. While this is good practice in most studies, it is especially important here for drawing strong conclusions.

### Other regressors may offer improvements

The principal difference between this study and those that came before it is the regressor. Because the auditory system is fundamentally nonlinear (viz., it responds with the same sign to both compression (positive) and rarefaction (negative) clicks, some sort of manipulation of the audio into an all-positive signal is needed. Previous studies have used the amplitude envelope (Aiken and Picton, 2008; Lalor and Foxe, 2010), spectrotemporal representations (Ding and Simon, 2009), and even dynamic higher-order features of speech (Di Liberto and Lalor, 2017).

Critically, the rectified speech audio used here is a broadband signal, which is what allows distinct ABR components at short latencies to be resolved in the derived response. There are many other transformations one could do, which will have important effects on the response waveform obtained. We piloted several (for example, “raising” the audio to be all-positive by adding it to its Hilbert amplitude envelope), but decided on the half-wave rectified audio due to its simplicity and the robustness of the responses it yielded. It is possible—likely, even—that there are better transformations. One shortcoming of our approach is that no distinct wave I was found, and all of waves I–V were smeared together. An improvement in the regressor is the most likely route to addressing this, and will be a focus of future work.

### Conclusions and future directions

Here we present and validate a method for determining the response of the auditory brainstem to continuous, naturally uttered, non-repeated speech. Speech processing involves a complex network that ranges from the earliest parts of the auditory pathway to auditory and association cortices. The techniques described here facilitate new neuroscience experiments by making it possible to measure activity across the auditory neuraxis while human subjects perform natural and engaging tasks. These paradigms will allow study of the subcortical effects of language learning and understanding, attention, multisensory integration, and many other cognitive processes.

## Acknowledgements

This work was funded by NIH grants R00DC014288 awarded to RKM and R01DC013260 awarded to AKCL. A preliminary version of this work was presented at the MidWinter Meeting of the Association for Research in Otolaryngology in February 2017. The authors wish to thank Susan McLaughlin and Tiffany Waddington for assistance with data collection.

